# Peptide-HLA II interaction prediction for post-translationally modified peptides

**DOI:** 10.64898/2026.08.07.743493

**Authors:** Alexandru Dumitrescu, Dani Korpela, Adrian M. Bebenek, Anna Ju, Griffin M. Lawrence, Karl R. Clauser, Jennifer G. Abelin, Martin Stražar, Harri Lähdesmäki, Daniel B. Graham, Ramnik J. Xavier

## Abstract

CD4^+^ T cells recognize peptides presented by human leukocyte antigen (HLA) II, implementing a fundamental mediation mechanism of the adaptive immune system. Although post-translational modifications (PTMs) alter immune responses, PTM-peptide-HLA interaction prediction remains challenging due to data scarcity resulting from substoichiometric levels of PTMs. To overcome this, we developed PepChem, a deep learning model utilizing novel, molecular-level peptide representations that enable predictions for sidechain modifications. Using monoallelic datasets that we reanalyze for PTMs of interest, we show accurate predictions on PTMs that were unseen during training. Furthermore, we introduce a novel training protocol that improves PTM-peptide generalization compared to conventional methods. We predict and experimentally validate citrullination-induced binding increase of rheumatoid arthritis (RA)-linked peptides to HLA II risk allele DRB1*04:01. This framework bridges the critical gap in PTM-aware immune recognition prediction, with immediate applications in autoimmunity, cancer, and infectious disease.

## 1. Introduction

The cellular arm of the adaptive immune response is mediated by T cells, which utilize germline rearranged antigen receptors to recognize diverse peptide antigens presented by HLA II on accessory cells. CD4^+^ T cells are restricted to peptides presented on HLA II and coordinate adaptive and innate immune responses customized to a wide variety of taxonomically diverse pathogens; however, CD4^+^ T cell reactivity to self antigens can elicit autoimmunity, which is further supported by genetic association of HLA II with autoimmunity and inflammatory diseases. The mechanisms that limit CD4^+^ T cell responses to self antigens involve central and peripheral tolerance, although there are key gaps in our understanding of tolerance failure. In central tolerance, developing T cells specific for self antigens are deleted by negative selection in the thymus. Self antigens that are unseen in the thymus but emerge in the periphery, however, can induce self reactive autoimmunity. In this regard, PTMs that occur only in peripheral tissues or during inflammation can be perceived as foreign antigens by CD4^+^ T cells (Doyle & Mamula, 2012). We sought to develop computational approaches to identify neoepitopes created by PTMs.

Obtaining experimentally generated PTM-peptide presentation data for HLA II, which is critical for training and evaluating machine learning (ML) models, remains a significant experimental challenge. The substoichiometric levels of PTMs pose a major problem for their identification without prior enrichment steps. Immunopeptidomics identifies peptides bound to HLA proteins without applying enzymatic digestion (no enzyme approach), which is essential for studying naturally presented peptides on the cell surface. The non-specific enzyme generated procedure, however, expands the search space dramatically, which is further amplified with each considered PTM. Previous efforts to identify PTM-containing peptides in the immunopeptidome have been made for HLA-I (Kacen et al., 2023), while our study uniquely introduces HLA II immunopeptidomics data containing PTM-peptides.

Current tools for predicting peptide binding to HLA II have proven remarkably successful in antigen discovery but remain constrained by their focus on the 20 canonical amino acids. Early approaches employed position weight matrices to identify amino acid enrichment patterns, yet their predictive power is limited by considering only the independent contributions of core residues to binding affinity (Nielsen et al., 2007). Subsequently, neural networks were introduced for peptide-HLA II prediction (Nielsen et al., 2008), evolving to incorporate additional biologically relevant information such as cellular localization (Graham et al., 2018), flanking amino acids relevant for cellular processing and presentation (Nilsson et al., 2023), and the peptide’s source protein (Strazar et al., 2023). Inspired by the success of protein language models in diverse biological domains (Elnaggar et al., 2022), deep contextualized latent representations have also been leveraged for peptide-HLA II binding prediction (Cheng et al., 2021). More recent exploratory efforts employing structural modeling (Abramson et al., 2024; Passaro et al., 2025) show promise in improving peptide-HLA (pHLA) binding core annotation but are currently outperformed by sequence-based methods in filtering non-binders (Ko et al., 2024). No prior ML models for peptide-HLA II interactions predictions can analyze post-translationally modified amino acid sequences.

To address these limitations, we propose novel molecular-level representations designed to bridge commonalities between PTMs and standard amino acids, thereby enhancing generalization. We denote our method PepChem, comprising ensembles of deep neural networks (NNs) trained on chemical representations of peptides that simultaneously predict the interaction strength between peptides and HLAs and detect their most likely binding regions.

Our method results in highly accurate prediction performance for both standard and PTM amino acids. We evaluate our model using reanalyzed HLA II immunopeptidomics containing PTMs, and present a novel training procedure that can incorporate PTM peptides in an unbiased manner.

## 2. Results

### 2.1. PepChem: a peptide-HLA II interaction prediction model for post-translationally modified peptides

#### Overview

PTMs, of which approximately 700 have been listed in UniProt (UniProt Consortium, 2025), alter the side chains of standard amino acids, which can significantly change peptide interactions with HLA II proteins. Different representations exist for the 20 *protein-coding* amino acids encoded by DNA, ranging from simple categorical variables (one-hot encoding) and substitution matrices (Henikoff & Henikoff, 1992) to embeddings from large protein language models (Lin et al., 2023; Elnaggar et al., 2022); however, they are limited by their finite vocabularies, and cannot accommodate arbitrary chemical side chain modifications. Here, we design a data-independent, molecular-level representation capable of capturing this expanded chemical space and enabling binding predictions for peptides that contain previously unseen PTMs. This representation, termed PepChem features, underpins a deep learning model, which simultaneously predicts binding affinity and presentation probability for any HLA II molecule and peptide pair.

The PepChem model is simultaneously trained on *in vitro* peptide-HLA II binding affinity (BA) data and peptide presentation from engineered antigen-presenting cells, often termed immunopeptidomics or eluted ligands (EL). This multi-task learning approach incentivizes shared representation learning between the two highly related tasks, resulting in improved BA performance as shown in our ablation studies. BA reflects the binding strength of a peptide to an HLA protein, reported in nanomolar concentrations, whereas EL annotations are binary indicators that capture both binding and additional aspects of antigen presentation, such as proteolytic processing or the presence of cleavage sites. Our model simultaneously predicts peptide-HLA II BA and EL probabilities based on the peptide sequence, the HLA II protein encoded by a subset of groove residues as described in (Nilsson et al., 2023), and optional contextual information from the peptide’s source protein (**Figure 1a**). All amino acids from the peptide, HLA II, and context are first encoded using the PepChem features, transformed using a learnable linear map, and then passed to an ensemble of NNs.

**Figure 1:**
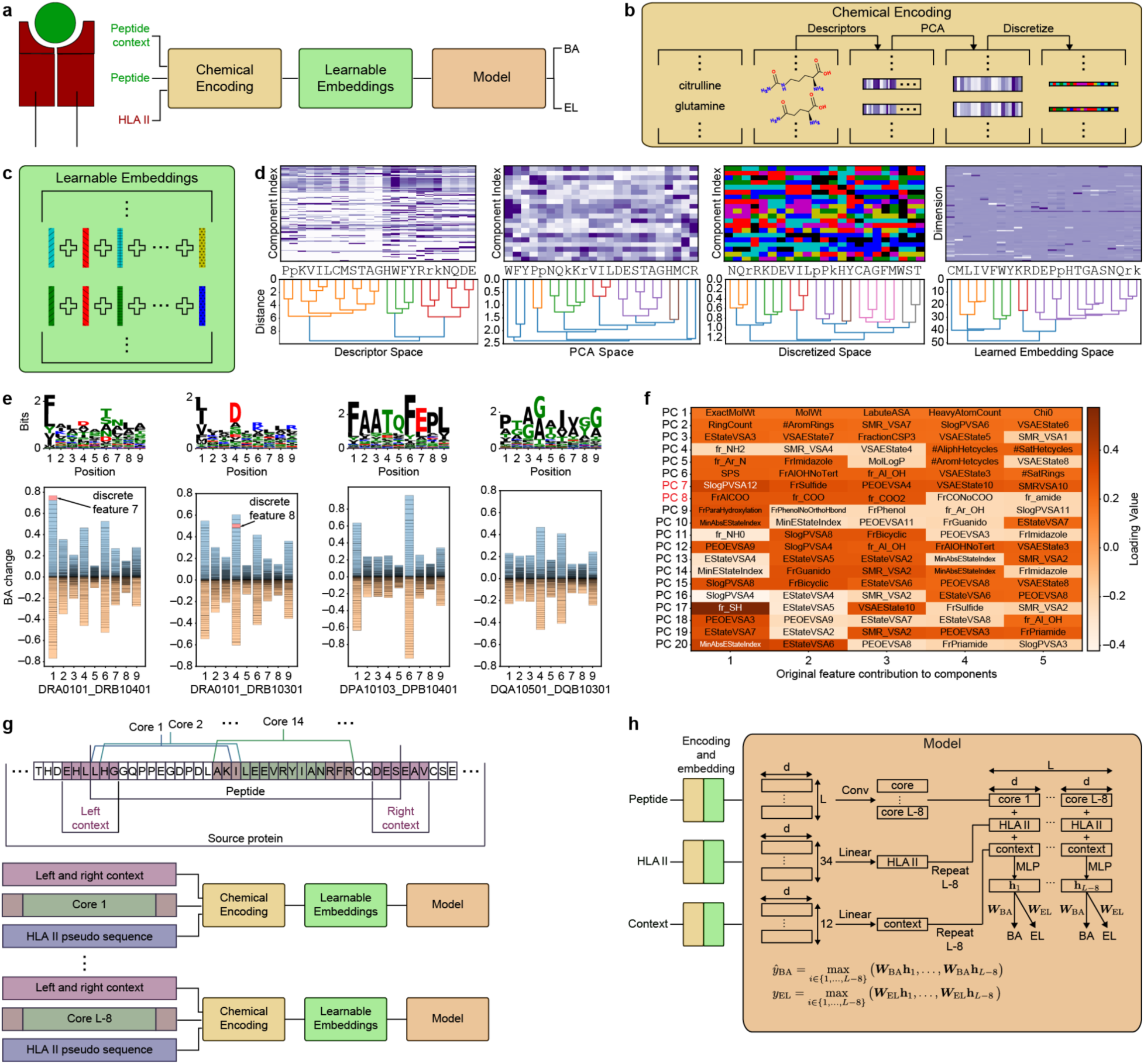
Molecular-level representations for PTM-peptide-HLA II interaction predictions. **a** Pipeline depiction. Our model predicts BA and EL based on peptide, HLA II, and, optionally, peptide source protein context, utilizing chemical representations. **b** Illustration of amino acid sequence transformation. **c** The first input layer of our model aggregates each amino acid’s corresponding learnable vectors to form the final representation of an amino acid. **d** Visualization and clustering of amino acids in four embedding spaces utilized in Chemical Encoding and Learnable Embeddings modules. We depict three PTMs, along with protein-coding amino acids, denoted by lower case letters: citrulline (r), acetyllysine (k), and hydroxyproline (p). **e** *Upper*: extracted motifs based on nonamer cores of high BA peptides; *lower:* effect of every discrete chemical feature category at each core location to binding affinity for different HLA II alleles. **f** The principal components along with the raw chemical descriptors with the highest loadings (in absolute value). **g** Predictions are made based on nonamers and contextual information, which includes 12 amino acids from the source protein around the peptide termini. Source protein amino acids are optional. **h** Architecture of one NN in our ensemble. Multilayer perceptron (MLP) block depth may vary.

#### PepChem features

The PepChem representation leverages the molecular graph information of each amino acid and transforms these into continuous vector embeddings of chemical descriptors or “molecular fingerprints” using RDKit <u>(Bento et al. 2020)</u>. These encode 210 key properties such as hydrophobicity, molecular size, solvent accessibility, polarity, surface areas, hydrogen bonding potential, and charge. We use 128 of those features that have non-zero variance across the standard amino acids and refer to these values as raw features.

**Figure 1b** illustrates the conversion of an amino acid sequence into PepChem features. We first apply Principal Component Analysis (PCA) to the raw features, compressing them to a subset of principal components (PCs). We assume that small differences in continuous chemical representations will not significantly alter binding affinities. Hence, we discretize the individual PCs using Gaussian Mixture Models (GMMs), which group chemically-similar amino acids into discrete classes within each component. This yields a discretized, molecular-level representation of each residue (**Methods Section 4.1**). We refer to this feature space as discretized features or PepChem features.

We then apply principal component analysis (PCA) to the raw features, yielding *n* principal components (PCs), each of which is subsequently discretized into a component-specific number of classes. The final discretization step proved crucial for generalization to PTMs, as we show later in our ablation studies. In essence, the discretization groups the amino acids in *n* different ways, based on different chemical characteristics, summarized by our PC representations.

Every class of each discrete feature has an associated learnable representation. All amino acids, including PTMs, will be represented as a sum of their corresponding learnable vectors (**Figure 1c**). Notably, each vector used in a PTM representation’s summation corresponds, by construction, to at least one other amino acid in the standard set, and is unique only through the set of all 20 vectors.

In **Figure 1d**, we show all standard amino acids and three additional PTMs in three representation spaces (raw features, PC, and PepChem features) and the learned embedding representation. The three chemical spaces determine how amino acids should be grouped.

Subsequent training of the PepChem prediction model refines the learned embedding to optimize prediction of peptide-HLA II binding. For example, in the learned embedding space, hydrophobic amino acids group together, and glutamine groups with citrulline and acetyllysine.

The key advantage of the discrete PepChem features is their straightforward generalization to residues not seen during training. We incentivize the model to learn the impact of each discrete feature using a special dropout training protocol (named PepChem_indep) which, with some probability, represents a few amino acids in a peptide using only a single categorical feature (**Methods Section 4.1**)—the same technique also allows us to incorporate PTM EL data during training, which otherwise converges to a degenerate solution where all PTM-peptides are predicted as binders (**Section 2.4**). We can therefore predict which of these discrete features has the largest effect on BA at certain core locations (**Figure 1e**). As expected, the largest affinity effects are observed at core anchor locations (**Figure 1e**).

The model identifies that the largest BA changes result from categorical features that are derived from relevant chemical information (**Figure 1f**). For example, the seventh discretized PC feature at position 1 for DRB1*04:01 has the largest gain of BA (**Figure 1e**) and is, in part, determined by water/octanol partition coefficient (LogP), while PC feature 8, dependent on the counts of the amino acid’s carboxyl groups, has the third largest increase in BA for position 4 of DRB1*03:01. This analysis suggests that the model associates the BA changes with appropriate chemical properties conveyed by the descriptors, including acidity and hydrophobicity.

#### PepChem model

The binding affinity of a peptide to HLA II proteins is primarily determined by a contiguous subset of nine amino acids—the binding core—which directly interacts with the HLA binding groove. Identification of binding cores may pinpoint key immunogenic sequence features and has been addressed previously (Nilsson et al., 2023; Racle et al., 2023). To this end, we construct a large ensemble of NNs, each predicting BA and EL probabilities of all possible nonameric substrings of the peptide. Every NN in the ensemble separately receives all possible peptide cores, along with context and HLA II information (**Figure 1g**), and only the highest predicted BA and EL cores (respectively) are used for the loss computation (**Figure 1h, Methods Section 4.1**). In contrast to previous approaches, our framework increases the depth of individual networks by parallelizing their training, enabling greater representational capacity.

### 2.2 PepChem achieves highly competitive accuracy in standard amino acid pHLA presentation and binding prediction

We train our method on the dataset compiled by (Nilsson et al., 2023), which includes 125,985 peptide–HLA II pairs with quantitative BA measurements and 248,219 EL pairs. As there are no experimentally validated negative EL examples, each positive EL peptide is supplemented with ten additional non-overlapping segments from the same source protein. We refer to the combined BA and EL dataset as D*_net_*.

In all our benchmarks, we include two versions of our method, denoted as *PepChem* and *PepChem_indep*, where the former uses exclusively D*_net_* during training, while the latter contains half of the PTM EL dataset generated in this work (**Section 2.4**). We compare our method to MixMHC2pred (Racle et al., 2023) and NetMHCIIpan-4.3 (Nilsson et al., 2023), which utilize a similar, but shallow MLP ensemble, BERTMHC (Cheng et al., 2021), which utilizes a large protein language model, and CAPTAn_core (Strazar et al., 2023). To compare against the latter, we re-train our method on the same EL data splits from (Strazar et al., 2023).

#### Eluted ligand evaluation

The EL dataset from (Strazar et al., 2023), D*_CAPTAn_*, is used as our independent EL benchmark when comparing against all prior methods except CAPTAn_core. Comparing against CAPTAn_core is done separately, using the train, validation, and test splits defined in (Strazar et al., 2023). When using D*_CAPTAn_* as an independent test dataset, we remove all peptide-HLA pairs present in D*_net_* to obtain 161,741 presented peptides and 1.04 million protein segments annotated as negatives.

Using macro average AuROC across all alleles to assess prediction performance, PepChem model outperforms all other methods, with higher scores on the majority of alleles when tested on all data (**Figure 2a** *left*, 0.801 compared to next best model BERTMHC 0.770, *p* ≈ 1.7 ⋅ 10^-10^, *N* = 78 alleles). On sequences of the same length as used in training, it is superior against all except BERTMHC (**Figure 2a** *right* 0.847 compared to BERTMHC 0.843, not significant). The additional NN capacity proves crucial for peptide presentation performance, as our method and BERTMHC significantly surpass the shallow NNs (Racle et al., 2023; Nilsson et al., 2023) (**Figure 2a** *right*). BERTMHC falls behind on sequence lengths outside training data (**Figure 2a** *left*), likely because it uses the entire peptide for its prediction, whereas our method is restricted to nonamer cores and limited contextual information. Similarly, the PepChem model significantly outperforms CAPTAn_core (0.877 compared to 0.839, *p* ≈ 3.3 ⋅ 10^-24^, *N* = 78 alleles). Finally, we observe a consistent increase in performance when including source protein contextual information (**Figure 2b**), which informs the model of cleavage site presence.

**Figure 2:**
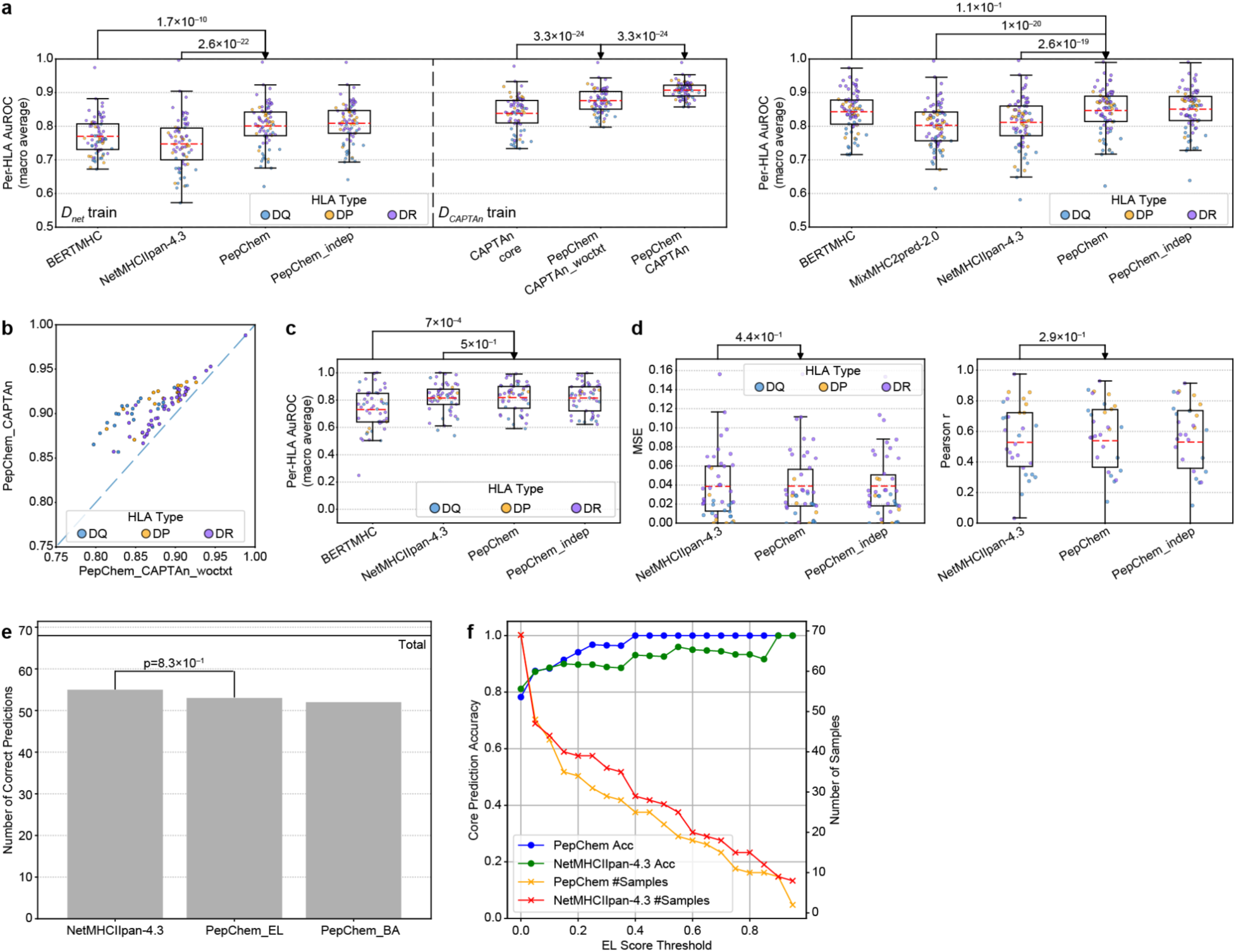
PepChem achieves highly accurate standard amino acid peptide-HLA II interaction predictions. **a** *Left*: testing peptides of the same length as the training; *right:* testing all peptide sequences. Performance of individual alleles (scatter points) and macro AuROC averages (red lines inside the box plots) are reported. The test measures the separation between positive EL peptides in D*_CAPTAn_* data and non-overlapping segments from their source protein, which are considered negatives. **b** Comparison between our model tested with (PepChem_CAPTAn) and without 6 amino acid context (PepChem_CAPTAn_woctxt) from the source protein. **c** Epitope benchmark with known HLA II specificity. This test considers epitopes known to contact HLA II proteins, along with all other segments of equal length from their source proteins, annotated as negatives. We predict BA and compute the separability of the epitopes from the created set of negatives. Each dot represents one HLA II, and the red lines inside the box plots depict the macro AuROC average performance. **d** IEDB benchmark. We report mean squared error (MSE, *left*) and linear correlation coefficients (*right*) for each allele, and depict macro averages as red lines inside the box plots. **e** Core prediction performance. **f** Core prediction accuracy over increasing peptide-HLA II probabilities and remaining number of samples for each EL score threshold. *P values are computed using one-tailed binomial tests for all results except core predictions (panel **e**), which use Fisher’s exact test*.

#### Binding affinity evaluation

We assess standard amino acid BA prediction performance using two datasets. First, we use the benchmark data collected by (Nilsson et al., 2023), resulting in 666 epitopes with known HLA restrictions and CD4^+^ binding T cells after removing duplicates, peptide-HLA II pairs present in the BA training dataset, and HLA II alleles outside of the training data, which we denote D*_NetBenc_*_ℎ_. As noted in (Nilsson et al., 2023), these epitopes are individually selected and are not necessarily presentable peptides. Each epitope, along with all equal-length disjoint segments from its source protein (marked as negatives), are used to compute AuROC scores. The second dataset, D*_IEDB_*, contains continuous BA measurements for peptides-HLA II pairs from IEDB (Vita et al., 2025). We only consider peptides that have no nonamer in common with any peptide from our training set D*_net_*, comprising 2,630 peptides.

The prediction performance of PepChem on BA data is highly competitive with prior methods, although it exhibits fewer gains from increased NN capacity (**Figure 2c**,**d**). We postulate that BA is often determined by four anchor positions within a binding core. To explain these cases, shallow networks likely have sufficient capacity. Although more complex behaviour has been noted, including potential multiple binding cores within the same peptide (Tong et al., 2006), we observe overfitting on the BA task. This suggests that some level of noise in the BA annotations is present, prohibiting performance gains of deeper networks.

Lastly, PepChem features obtain the same performance as categorical representations (one-hot encoding) and substitution matrices (Henikoff & Henikoff, 1992) on standard amino acid data (**not shown**). In addition, jointly training the model on the EL and BA predictions results in improved BA performance (**not shown**).

#### Core predictions

To assess core prediction performance, we compared predicted cores to those observed in experimentally solved peptide-HLA II structures. We used benchmark datasets from (Mikhaylov et al., 2024) and (Ko et al., 2024), filtered to retain only human MHCs and resulting in 69 peptide-HLA II pairs across 25 different alleles. All NNs detect core locations based on their highest BA or EL score across the peptide’s nonamers, and the ensemble’s final core prediction is determined by majority voting.

In **Figure 2e**, we compare the number of correct cores detected using PepChem’s EL predicted scores to that of NetMHCIIpan-4.3 predicted cores. In **Figure 2f**, we show that EL confidence estimated by PepChem is the most reliable measure of core accuracy for PepChem, compared to percentage of votes or BA scores (**Supplementary Figure 3**).

In summary, PepChem features and simultaneous usage of EL and BA data provide the model with molecular-level information and augmented training dataset size. When limited to canonical amino acids, this translates to competitive performance in both prediction tasks as well as binding core inference.

### 2.3 Immunopeptidomics identifies HLA II allele-specific binding preferences for peptides with PTMs

To assess presentation performance and enable generalization on peptides containing PTMs, we generated an EL dataset including these PTMs by reanalyzing publicly available monoallelic HLA II data using a tailored search protocol (**Methods Section 4.2**). The PTMs considered are citrulline, acetyllysine, hydroxyproline, cystine, pyroglutamic acid, oxidized methionine, and asparagine deamidation, with observed presentation annotations for 83 different HLA II heterodimers.

Our reanalysis initially identified 132,713 unfiltered peptides containing one or more PTMs in addition to canonical amino acid peptides. Because global false discovery rate (FDR) estimation underestimates the true error rate for rare modifications, we applied group-level FDR so the modified and unmodified peptides are subject to a strict 1% FDR within each category (**Methods Section 4.2**). We then filtered these identifications further using quality metrics including strict mass tolerances and sequence-coverage requirements to ensure the accurate identification of class II HLA peptide ligands with various PTMs (**Figure 3a**, **Methods Section 4.2**). We paid particular attention to the ambiguity between citrullination of arginine and deamidation of asparagine, a well-documented source of PTM misassignment. These controls ensure that the modified ligands used downstream reflect confidently localized PTMs.

**Figure 3:**
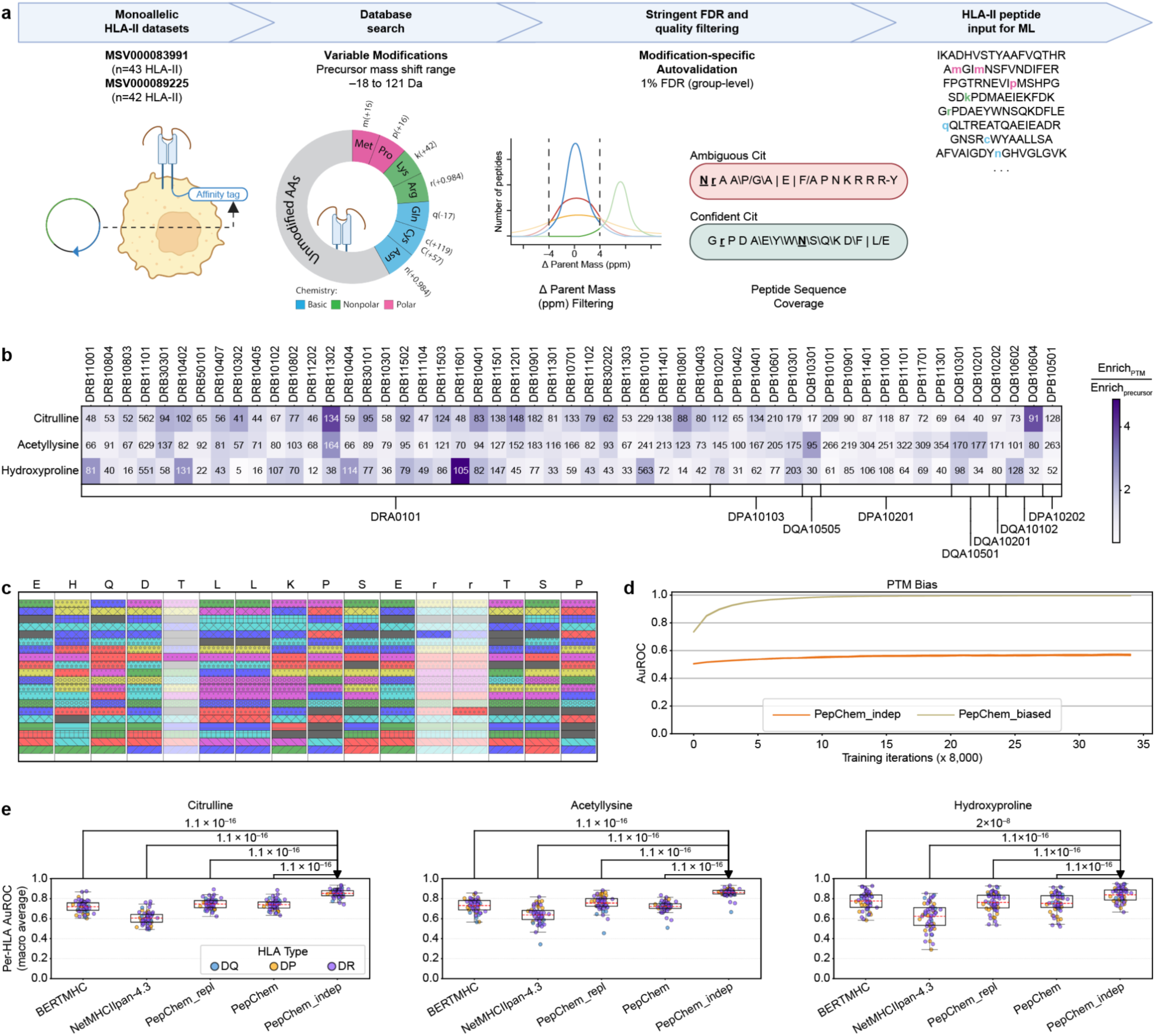
PepChem achieves highly accurate PTM amino acid peptide-HLA II interaction prediction. **a** PTM data extraction pipeline. **b** PTM peptide data statistics. Purple shades represent the PTM enrichment divided by the protein-coding precursor, for each allele. The numbers within each box denote the total number of PTMs. **c** Visualization of the feature dropout. **d** Bias test, measuring how likely a random PTM mutation is to increase the predicted EL presentation probability. We measure the bias throughout training to determine whether each method has an uncontrollably increasing bias. **e** Separation test between presented peptides containing PTMs and non-presented protein segments containing only standard amino acids. *P values are computed using one-tailed binomial tests for PTM-peptide AuROC performance (panel **e**)*.

Overall, following this stringent identification and disambiguation pipeline, we observed the presentation of peptides bearing the modifications above across all 83 HLA II heterodimers, providing a high-confidence foundation for modeling how PTMs influence allele-specific ligand presentation.

### 2.4 Chemical descriptors enable accurate modeling of post-translationally modified peptides binding to HLA II

After mass tolerance and sequence ambiguity filters, we remove duplicates and restrict the considered dataset to peptides with HLA II annotations present in D*_net_*, resulting in 53 distinct heterodimers. We consider three PTMs of interest for PTM training and validation analysis, resulting in 12,280 peptides containing at least one citrulline, acetyllysine, or hydroxyproline (**Figure 3b**). We refer to this filtered dataset as D*_PTM_*_–*EL*_. Finally, we split the PTM-containing HLA II peptides into two equally-sized subsets, training on one and predicting on the other. This data is not included in the validation set, nor does it affect early stopping.

To incorporate the newly derived PTM EL data, we develop a new training method, PepChem_indep, that does not require a PTM negative set. We design a special, feature-level dropout that encodes PTMs and random subsets of the standard amino acids using only a single, randomly chosen discrete feature (**Figure 3c, Methods Section 4.1**). Note that, by construction, all categorical features of PTMs overlap with at least one standard amino acid.

This dropout technique forces the model to learn binding properties from features themselves and not memorize their fixed combinations. Effectively, this avoids degenerate models where, for example, any peptide containing a PTM will be predicted as positive.

We first test whether peptides containing PTMs are systematically assigned different presentation probabilities than those with standard amino acids, by synthetically introducing PTM point mutations into random peptides and predicting presentation scores for both forms. Crucially, the proposed PepChem_indep method does not suffer from a large bias and eventually stabilizes over the training (**Figure 3d**). Conversely, we show that naively adding PTM data in the training set (denoted by PepChem_biased in **Figure 3d**) grows the model’s PTM-peptide probability uncontrollably: since no negative post-translationally modified peptides are present, the degenerate solution of assigning maximal probability to any PTM-containing peptide successfully optimizes the objective function.

We next measure the models’ ability to distinguish standard amino acid negative peptides present in (Strazar et al., 2023) from the set of positive PTM-peptides in D*_PTM_*_–*EL*_. To benchmark against prior methods, we replace PTMs with their closest corresponding standard amino acid in terms of Euclidean distance in the PC embedding space. PepChem generalizes well on unseen PTMs and results in highly competitive performance relative to prior methods (**Figure 3e**), with similar performance when replacing PTMs with their closest standard amino acid (PepChem_repl in **Figure 3e**). When including this data in training (cross-validated using two roughly equal folds) using our proposed dropout technique, we achieve significant performance gains across all considered PTMs, reaching 0.850 macro AuROC compared to next best model BERMHC 0.719 (*p* ≈ 1.1 ⋅ 10^-16^, *N* = 53 alleles) for citrullinated peptides, 0.858 compared to 0.730 (*p* ≈ 1.1 ⋅ 10^-16^, *N* = 53 alleles) for acetyllysine peptides, and 0.835 compared to 0.774 (*p* ≈ 2 ⋅ 10^-8^, *N* = 53 alleles) for peptides containing hydroxyproline (PepChem_indep compared to BERTMHC in **Figure 3e**).

Lastly, we also consider an alternative way of including this data in training, supplementing the positive PTM-peptides with synthetic negative PTM-peptides, by mutating precursor amino acids of negative peptides based on probabilities extracted from, which we denote PepChem_negPTMs. PepChem_negPTMs successfully mitigates the PTM bias and achieves good performance in separating EL PTM-peptide positives from standard negatives (**Extended Data Figure 1a,b**), but severely worsens the BA predictions on citrulline peptides (**Extended Data Figure 1c-e**). We highlight two major challenges for this method. First, our mutation probabilities used to create negative PTM-peptides are formed independently of the peptide sequence context, and thus may result in unnatural PTM-peptides. The model might exploit this, learning to identify natural PTM mutation sites to the detriment of peptide presentation characteristics. Works like (Peng et al., 2025; Wen et al., 2025) study PTM site prediction for a restricted set of PTMs (not including citrulline) and may, in principle, be employed to circumvent the positive and negative PTM-peptide distribution discrepancy. Second, some peptides may become binders as a result of the introduced synthetic mutations, inevitably introducing unknown amounts of false negatives in the training set. Conversely, our proposed dropout technique mitigates both of these challenges, since no artificial negative PTM-peptides are needed to achieve an unbiased model.

### 2.5 PepChem accurately models point mutation-induced gain and loss of function

We measure the effect of point mutations by computing BA differences between paired peptides of the same length differing by one amino acid (**Figure 4a**). We denote an increased BA resulting from a certain point mutation as gain-of-function and, conversely, a loss-of-function when the BA is lowered as a result of that mutation.

**Figure 4:**
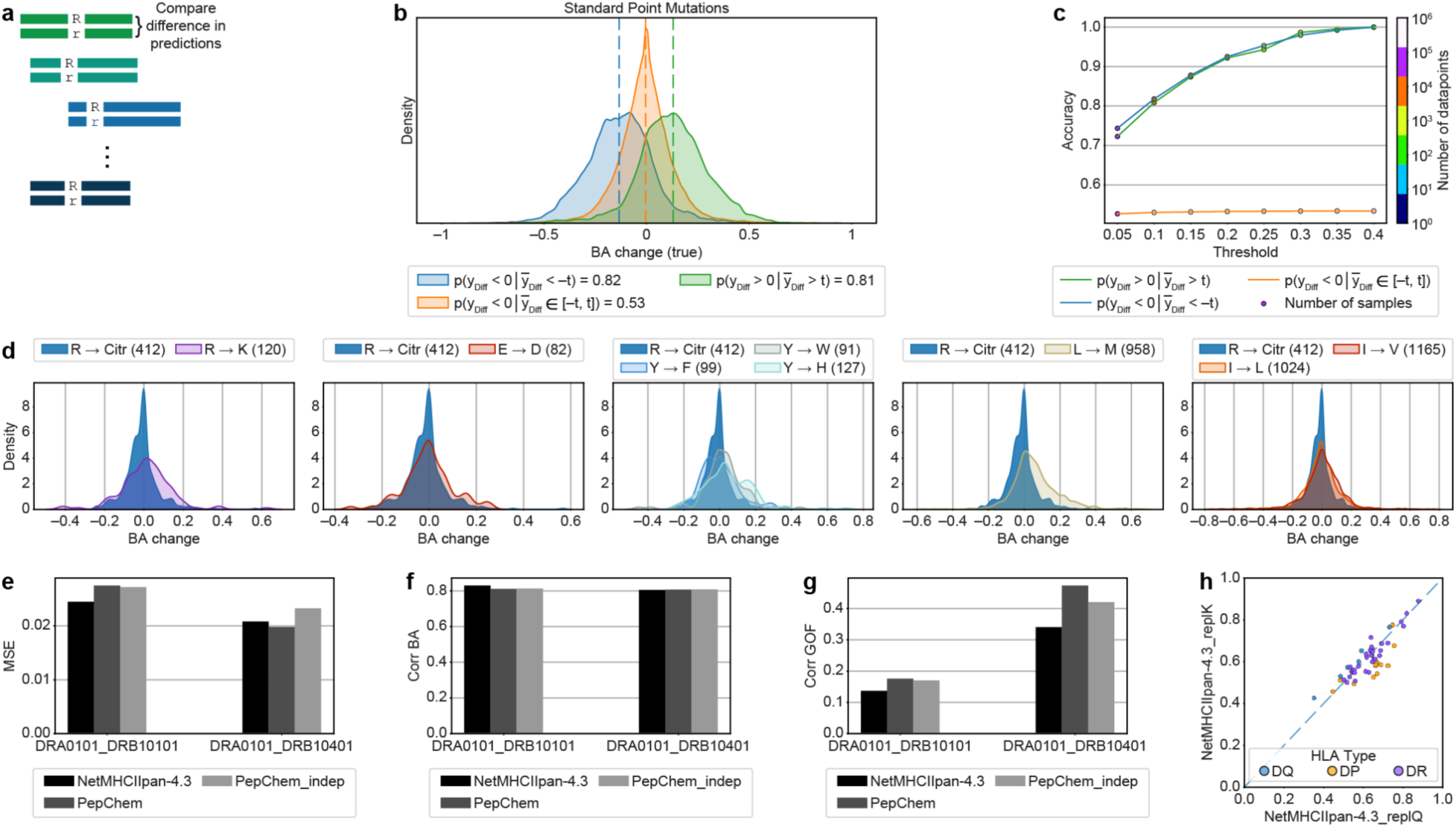
PepChem detects point mutation-induced peptide-HLA II BA changes. **a** We consider paired peptide sequences, where the two peptides differ by one amino acid and have BA annotations for the same HLA II. **b** Threshold-based separation into three classes of peptide pairs with standard point mutations. We classify all pairs of peptides into one of three classes using a threshold of *t* = 0.1. The distributions of the experimentally annotated BA differences for pairs in each predicted class are depicted. **c** Accuracy of the classes with increasing threshold *t*. **d** Distributions of BA changes resulting from citrullination, when compared to standard mutations with functionally similar residues (BLOSUM62 similarity scores ≥ 2). **e** Mean squared error between predicted and true BA on citrullinated peptides. **f** Linear correlation coefficients between predicted and true BA values for citrullinated peptides. **g** Linear correlation coefficients for the predicted BA difference between arginine and citrulline form peptides. **h** Separation performance of citrulline-containing EL peptides and a set of negative, standard amino acid peptides for various alleles.

#### Point mutations of standard amino acids

We first measure our method’s ability to capture BA changes resulting from standard amino acid point mutations. We use the data curated by (Nilsson et al., 2023), and extract all peptide pairs differing by exactly one amino acid and having experimentally measured BA on the same HLA II. We assess the performance on the held-out validation set.

We classify peptide pairs based on their predicted BA differences and varying thresholds into three classes: *i*) gain-of-function, *ii*) loss-of-function, and *iii*) no change. We predict a peptide pair as one of the first two classes when the BA differences are greater or lower, respectively, than the threshold *t* or −*t*, and consider a pair as having no change when the absolute value of the BA difference is lower than *t*. The true class is determined by the experimentally annotated BA difference.

In **Figure 4b**, the predicted BA changes are already well separated even for a small threshold of *t* = 0.1, while in **Figure 4c**, the precision converges to near-perfect for large enough thresholds *t*.

#### Citrullination

We evaluated 216 annotated peptide pairs consisting of citrullinated and arginine-form sequences reported in (Sidney et al., 2017), each with experimentally measured binding affinities (in nM) for DRB1*01:01 and DRB1*04:01. This dataset exhibits very small citrullination-induced BA changes, even when compared to those caused by functionally similar point mutations in D*_net_* (**Figure 4d**), which poses challenges to the reliable assessment of gain-of-function predictions. Citrullination modifies critical biochemical features (such as side chain charge) known to influence protein interactions, making it unlikely to generally incur smaller BA changes compared to functionally similar standard residue mutations. Thus, the unusually small impact of citrullination likely stems from the biased selection of peptide pairs (vimentin and collagen peptides).

We measure the mean squared error and linear correlation coefficients between predicted and annotated BA values, and gain-of-function correlations for our method and NetMHCIIpan-4.3. As the latter does not support non-standard residues, we substitute citrulline with glutamine, which yields the highest gain-of-function performance. Overall, both models deliver comparable binding affinity predictions: NetMHCIIpan-4.3 performs slightly better on MSE and BA correlation for one of the two alleles (**Figure 4e,f**), and the PepChem model has a small advantage on gain-of-function performance (**Figure 4g**). Notably, the protein-coding replacement (glutamine) of NetMHCIIpan-4.3 was chosen on the same data used to report the test performance, which contains only two alleles. In **Figure 4h**, we use the reanalyzed PTM-peptide presentation data D*_PTM_*_–*EL*_ to show that the optimal standard amino acid replacement is HLA II-dependent, with lysine sometimes yielding better performance than glutamine based on AuROC performance of citrullinated EL peptides, suggesting that the reported NetMHCIIpan-4.3 performance obtained in **Figure 4e-g** may be overestimated.

Lastly, besides enabling the usage of PTM EL data during training (**Section 2.4**), the discretization step proved crucial on the BA generalization performance to PTMs. The discrete descriptors (PepChem features) input representations achieve superior performance to continuous descriptors (PCs) when tested on peptide BA predictions for citrullinated peptides.

### 2.6 PTMs confer increased binding of peptides to HLA II alleles linked to autoimmune diseases

We determine the HLA II alleles most susceptible to gain of function upon citrullination, lysine acetylation, and proline hydroxylation and identify the responsible core positions. Crucially, our chemical representation framework naturally extends core prediction to peptides with PTMs, unlike finite-vocabulary models. We use the data obtained by (Strazar et al., 2023), D*_CAPTAn_*, and select all positive peptides (presented by at least one HLA II allele) that contain between one and eight precursor standard amino acids (arginine, lysine, or proline). For a peptide containing n precursor standard amino acids, we enumerate all its 2*^n^* − 1 mutated versions and compare their predicted BA to the original peptide, using the 45 HLA II alleles with the most BA annotated peptides.

For each HLA II allele, we select the largest 1,000 citrullination-induced gain-(**Figure 5a**) and loss-of-function (**Figure 5b**) peptides, depicting the distribution of the BA changes. In **Figure 5a**, we find a perfect separation between rheumatoid arthritis (RA)-susceptible and RA-protective HLA II DR alleles, based on the HLA II associations to anti-citrullinated protein antibodies-positive (ACPA^+^) RA patients reported in (Kampstra et al., 2017). Notably, citrullination incurs the smallest gain of function on protective allele DRB1*13:01, while DRB1*04:01 is the third most susceptible to citrullination-induced gain of function (**Figure 5a**).

**Figure 5:**
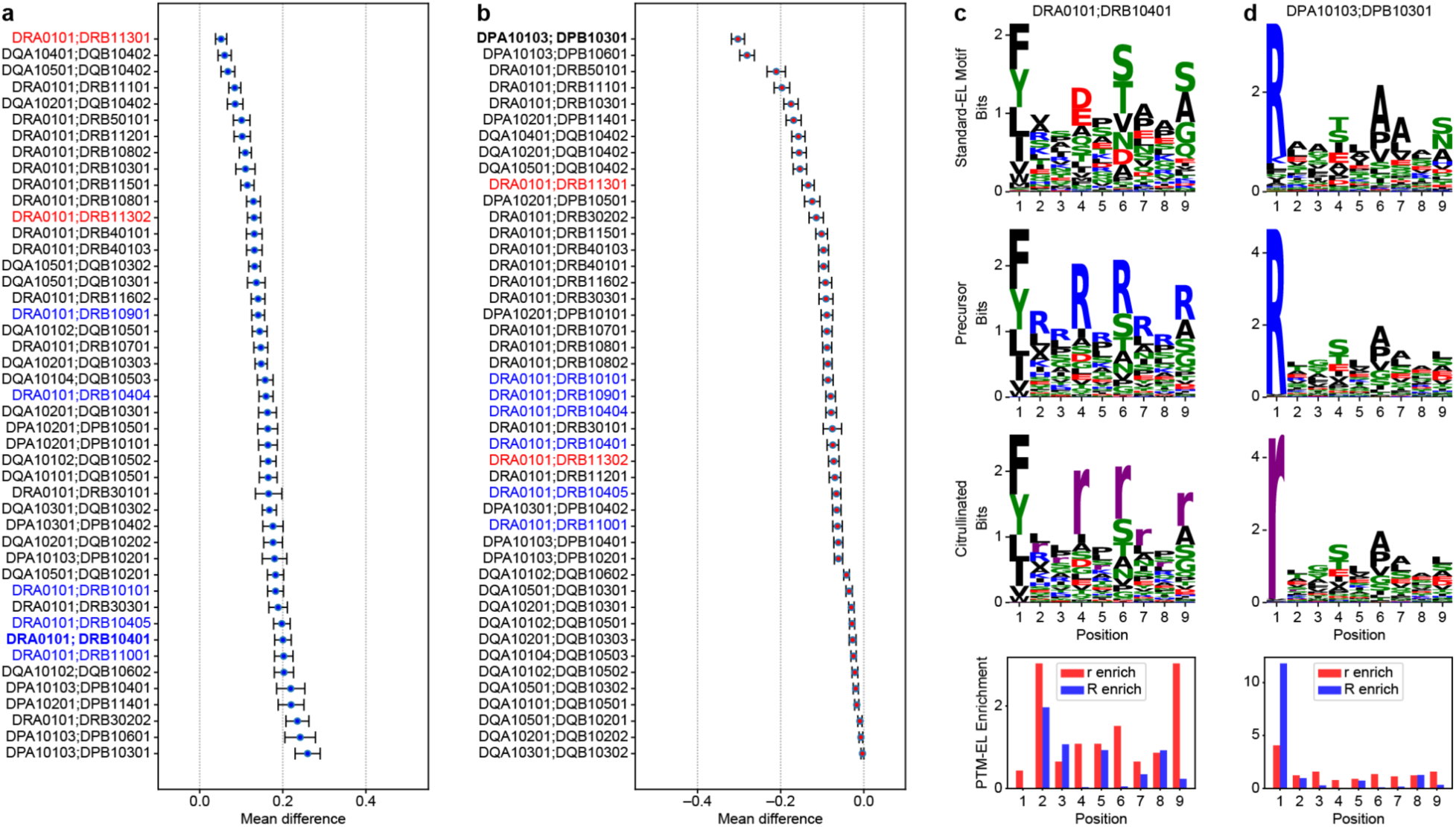
PepChem showcases preferential citrulline-induced gain and loss of function in peptide-HLA II interactions. **a** BA change distribution of 1,000 peptide pairs resulting in the largest gain of function from citrullination. Blue and red DR alleles are susceptible and protective RA alleles, respectively. **b** BA change distribution of 1,000 peptide pairs resulting in the largest loss of function from citrullination. Blue and red DR alleles are susceptible and protective RA alleles, respectively. **c** Gain-of-function analysis for DRB1*04:01. From top to bottom: *i*) DRB1*04:01 core extracted from the EL validation data of D*_net_*; *ii*) precursor motif that, upon citrullination, incurs the largest gain of function; *iii)* resulting modified motif that incurs a large gain of function upon citrullination; and *iv*) increased frequencies of citrulline and arginine at various positions based on the D*_PTM_*_–*EL*_ data, for the high gain- and loss-of-function alleles. **d** Loss-of-function analysis for DPA01*01:03;DPB1*03:01.

For DRB1*04:01, the model predicts that citrullination of arginine at positions 4, 6, and 9 results in the largest gain of binding affinity (**Figure 5c**). This may be due to absence of positively charged residues at these sites (**Figure 5c**, Standard-EL Motif). In contrast, as citrulline is not hydrophobic, citrullination at position 1 does not predict a substantial gain of function. This is corroborated by the enrichment patterns observed in the PTM EL dataset, where citrulline shows moderate to strong enrichment at positions 4, 6, and 9, accompanied by a corresponding depletion of arginine (**Figure 5c**, PTM-EL enrichment). Conversely, the core of DPA01*01:03;DPB1*03:01 (**Figure 5d**, Standard-EL Motif) contains an enriched arginine at position 1, which results in a loss of function when citrullinated. Based on the PTM EL data, position 1 of the core has a much larger arginine enrichment compared to citrulline.

Lysine acetylation had the largest predicted effect on BA (**Extended Data Figure 2a,b**), most notably with DQA1*05:01;DQB1*03:01, in agreement with our enrichment analysis flagging acetyllysine at positions 3, 4 and 6. (**Extended Data Figure 2c,** PTM-EL enrichment). Within DRB5*01:01, lysine acetylation at position 9 results in a loss-of-function prediction, likely due to the elimination of positive charge, a prediction that is corroborated by the PTM EL data (**Extended Data Figure 2d,** PTM-EL enrichment).

Finally, our approach predicts that proline hydroxylation yields minimal changes in binding affinity (**Extended Data Figure 3a,b**). The gain of function anticipated for DRB3*02:02 was not substantiated by PTM EL data (**Extended Data Figure 3c**, PTM-EL enrichment), likely due to limited data availability. In contrast, the predicted proline hydroxylation in **Extended Data Figure 3d** (PTM-EL enrichment) is corroborated by a stronger proline preference at position 6 compared to hydroxyproline.

### 2.7 Citrullination incurs higher affinity of RA patient synovial fluid peptides to DRB1*04:01

Citrullination is a well-established hallmark linked to RA, and the allele HLA-DRB1*04:01 is strongly associated with increased disease susceptibility (Weyand et al., 1992). We analyze 515 previously identified citrullinated peptides in the synovial fluid of RA patients by (Tutturen et al., 2014) and (Wang et al., 2016), and determine whether citrullination could contribute to immune activation through enhanced HLA II presentation by DRB1*04:01.

We ranked peptides based on the change in predicted binding affinity from the unmodified arginine form to the citrullinated form. **Figure 6a** highlights five citrullination events predicted to incur large increases in DRB1*04:01 binding affinity. For these pronounced gain-of-function peptides, we performed a dose titration to measure binding of fluorescently-labelled peptides to DRB1*04:01 immobilized on microbeads (**Method Section 4.3**). We show our model’s mean and standard deviation predictions over different cores in **Figure 6b** and the resulting titration experiment mean fluorescent intensity (MFI) over peptide concentration in **Figure 6c**. The two highest-confidence citrullination-induced gain-of-function predictions on peptide pairs 1 and 3 (AEFTGrHDAHLNGK and SQLVYQSrRGPLVK) showed substantial increases in binding upon substituting arginine with citrulline at position 4 of the predicted binding core. Peptide pairs 2 and 5 exhibited more modest gains, involving arginine-to-citrulline substitutions at positions 6 and 9, respectively, suggesting that DRB1*04:01 may tolerate or even slightly increase BA at more than one anchor location within the cores. By contrast, peptide pair 4, our lowest-confidence prediction, showed a slight loss of function, with the two peptide forms having distinct predicted binding cores.

**Figure 6:**
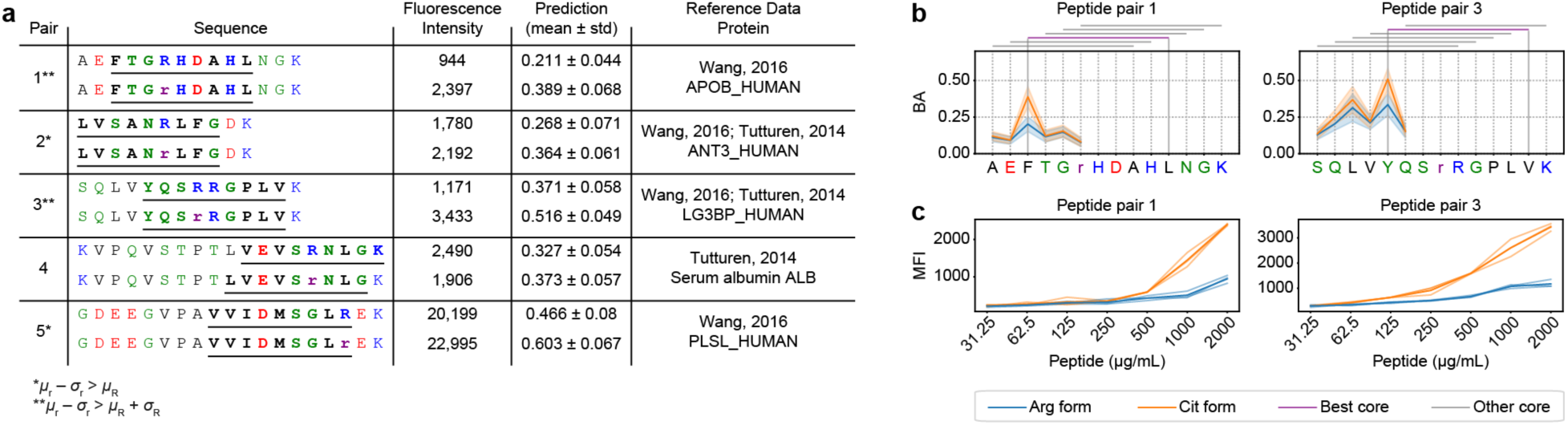
Peptides from RA patient synovial fluid result in a gain of function to RA risk allele when citrullinated. **a** Table containing peptide pairs with large gain-of-function predictions from arginine to citrulline form. Pair indices with one or two asterisks (*, **) correspond, respectively, to high and very high gain-of-function confidence predicted by our NN ensemble. The predicted cores in each sequence are underlined and bolded. The fluorescence intensity values in this table correspond to mean fluorescence intensity of FITC-labelled peptide bound to recombinant DRB1*04:01/DRA heterodimers on microbeads as measured by flow cytometry. The mean fluorescent intensity (MFI) values correspond to the largest peptide concentration (2,000*μ*g/mL) in our titration experimental validation (panel c). **b** Each nonamer’s predicted BA value is depicted at its starting location on the x axis, for the arginine (blue) and citrulline (orange) forms. The solid lines represent the mean prediction, and the shaded areas are one standard deviation over the NN ensemble. **c** Titration experimental validation of the RA peptide pairs selected based on the gain-of-function predictions, measured in MFI over peptide concentration. The indicated concentrations of FITC-labelled peptides were incubated with recombinant DRB1*04:01/DRA heterodimers attached to microbeads, followed by measurement of bead-associated fluorescence by flow cytometry.

These citrullinated peptides, identified *in vivo*, prioritized by PepChem, and validated to undergo gain of binding function, are promising starting points to improve our understanding of disease onset.

## 3. Discussion

We introduced PepChem, a novel machine learning approach that utilizes chemical representations of amino acids to enable peptide-HLA II interaction prediction for peptides with any side chain modification. Instead of fixed vocabularies, we represent each amino acid with continuous chemical features and discretize each dimension into a few representative classes.

To enhance generalizability, by construction, PTM categorical classes correspond to at least one other standard amino acid. Our approach results in highly accurate predictive performance on modified peptides unseen during training, validated on citrullinated peptides from (Sidney et al., 2017), and 12,280 new, modified eluted ligands generated in this work. The developed PepChem features achieve state-of-the-art performance on standard pHLA datasets, thus enabling generalization to PTMs without sacrificing performance on canonical amino acid data.

Machine learning-based structural prediction models have been extended and developed to include both proteins and small molecules. In particular, (Abramson et al., 2024) can predict structures for PTM proteins, but falls short on discriminating between weak and strong binders for p-HLA complexes (Ko et al., 2024). The extent to which structural models may aid binding affinity predictions on highly diverse modalities remains an open question. While some tasks, such as protein function prediction, can benefit from protein backbone structure alone (Gligorijević et al., 2021), protein-protein and peptide-protein interactions occur through their side chains, and therefore require structural predictions to accurately capture their orientations. Consequently, the related task of interaction prediction between the highly heterogeneous T cell receptors (TCRs) and pHLA complexes remains unsolved (Nielsen et al., 2024), and exhibits modest accuracy gains from structural predictions. Similarly, presented peptides may originate from taxonomically diverse organisms, resulting in a large degree of heterogeneity. Lastly, structural approaches suffer from significantly slower runtimes, typically employing a slow multiple sequence alignment step (Jumper et al., 2021; Abramson et al., 2024; Passaro et al., 2025), followed by multiple forward passes of larger NNs implementing iterative refinement (Jumper et al., 2021) or diffusion generative models (Abramson et al., 2024; Passaro et al., 2025) for a single biological complex query. This limits their applicability for large-scale *in vivo* peptidome analyses carried out in this work. Conversely, sequence-only peptide-HLA II interaction data is highly abundant, which enables the development of highly accurate and computationally efficient models, while retaining some structural interpretability through core predictions.

We additionally introduce a method of incorporating PTM EL data in training by applying a feature-level dropout. This circumvents the need for an artificial negative set, which is commonly employed to compensate for limitations of immunopeptidomics methods for standard amino acids, but introduces additional unpredictable biases in PTM contexts. The method relies on our discrete representation construction that assigns classes corresponding to one or more protein-coding amino acids to chemically-derived categorical features of PTMs. We test our novel training procedure by training and testing on the reanalyzed public HLA II data in a cross-validated manner, showing significant improvements. We envision the peptide encoding to be applicable to any classification task involving PTMs, while the dropout technique provides an unbiased way to include positive-only PTM data.

Our work fills an important gap in the analysis of PTMs in health and disease. DRB1*04:01 is a well-established RA-linked polymorphism (Weyand et al., 1992), and RA susceptibility is increased in the presence of anti-citrullinated antibodies (Macgregor et al., 1995). To this end, we showcased and experimentally validated peptides where citrullination induces large affinity increase of RA patient tissue-derived peptides to the RA risk allele DRB1*04:01. In addition, by predicting synthetic PTM mutation effects, we illustrate various anchor locations where citrullination of arginine, acetylation of lysine, and hydroxylation of proline may preferentially increase peptide-HLA II BA in certain HLA II alleles. Notably, we find that our model separates susceptible and protective DR alleles based on citrullination-induced gain of function, using HLA II associations reported for RA patients with anti-citrullinated protein antibodies (Kampstra et al., 2017). Our work established crucial groundwork for further investigation into the role of PTMs in autoimmune diseases, which are promising but yet unproven disease onset factors (Doyle & Mamula, 2012).

Our chemical descriptors are derived in a data-independent manner and are therefore suitable for any biological application considering residue side chain modifications (e.g. discovering antimicrobial peptides <u>(Wang et al. 2022)</u> or predicting peptide retention time <u>(Ma et al. 2018)</u>). An important unaddressed aspect is the effect of PTMs in TCR specificity, which was omitted given the persisting challenges of predicting TCR-pHLA complex interactions for arbitrary peptides-TCR pairs (Korpela et al., 2023; Bradley 2023; Nielsen et al., 2024). Nonetheless, future extensions of our work may also include TCRs, although PTM-pHLA-TCR analysis will likely require extensive experimental efforts and a restricted context, acquiring and considering sufficient data for a restricted subset of epitopes. As mass spectrometers and analytical methods are beginning to enable routine detection of post-translationally modified peptides (Kessler et al., 2025), future ML models will benefit from the ability of PepChem to represent common and novel PTMs and seamlessly incorporate the resulting data.

## 4. Methods

### 4.1 PepChem method

We represent standard and PTM amino acids using smiles encodings and obtain dense molecular representations using RDKit <u>(Bento et al. 2020)</u>, which we downscale and then discretize, assuming the compressed space follows a gaussian mixture. We ensure each PTM discrete feature also corresponds to at least one other standard amino acid.

Our model consists of a large ensemble of neural networks, similar to the ones developed in (Nilsson et al., 2023; Racle et al., 2023). In contrast to prior methods, we enable the usage of arbitrary depth, using a modern PyTorch implementation, which allows us to train multiple NNs in parallel, within the same GPU.

Compared to (Nilsson et al., 2023; Racle et al., 2023), we obtain better EL performance, since our higher capacity NNs can better learn the more complex relationships between sequence and peptide processing and presentation. However, deeper architectures are prone to overfitting and the BA data is noisy, which initially prohibited successful multi-task learning between the EL and BA objectives. To enable optimal transfer of shared patterns across the two highly related tasks, we re-expressed the continuous optimization objective through a binary one, which can be proven to converge to the same solution on optimality, while having a slower convergence curve.

Lastly, an important advantage of our representation is the ability of incorporating PTM EL data in training, which normally consists only of positive examples. Our method does not need any artificial negative datapoints, which we observe to significantly hinder the resulting PTM embeddings (significantly worsens the BA predictions of citrullination, **Extended data Figure 1c-e**, PepChem_negPTMs). Instead, we can incorporate only the additional positive samples by using the feature-level dropout that also eliminates the convergence to degenerate solutions.

### 4.2 Immunopeptidomics EL PTM data

Publicly available monoallelic HLA class II immunopeptidomics datasets were reanalysed to identify peptides carrying post-translational modifications of interest. Tandem mass spectra were processed using a standard proteomics database-search workflow against a human reference proteome supplemented with common contaminants. Searches allowed a predefined set of biologically and experimentally relevant peptide modifications and were performed without enzyme-specific cleavage constraints.

Peptide-spectrum matches were accepted using target–decoy-based false-discovery control and subsequently subjected to additional quality-control procedures based on mass accuracy, spectral evidence, peptide properties, and modification-site interpretability. Modified and unmodified peptide identifications were evaluated separately to account for differences in their respective search spaces and expected error profiles. High-confidence spectrum-level identifications were consolidated into distinct peptide-modification states, and peptide signal intensities were used as relative abundance estimates. Additional conservative filters removed identifications for which closely related modification assignments could not be distinguished reliably from the available fragmentation evidence.

### 4.3 Experimental validation PTM data

Fluorophore-labelled predicted peptides were obtained commercially and evaluated for binding to recombinant HLA class II molecules. Peptide binding reactions used recombinant biotinylated MHCII heterodimers and a concentration series of labelled peptides under mildly acidic, detergent-containing conditions designed to support peptide exchange. Reactions were protected from light and incubated to allow complexes to reach equilibrium.

Following incubation, peptide–MHCII complexes were captured on streptavidin-coated magnetic beads. Bead-associated MHCII was detected with a fluorescent anti-HLA-DR antibody and quantified by flow cytometry. Samples were handled under light-protective conditions, washed before acquisition, and analysed using standard cytometry-analysis software. Fluorescent peptide signal associated with captured MHCII complexes was used to assess relative peptide binding across the titration series.

Recombinant HLA-DR heterodimers representing the alleles examined in this study were produced in a mammalian expression system using an Fc-mediated heterodimerization strategy. Expression constructs encoded soluble extracellular HLA chains, peptide-loading elements, and tags enabling enzymatic biotinylation and affinity capture. Following transient co-expression, secreted material was clarified and purified by sequential affinity and size-exclusion chromatography. Purified complexes were biotinylated, buffer-exchanged, concentrated, and stored frozen until use.

## Supplementary information

Supplementary tables, derivations of machine learning methods in PepChem, and training parameters are described in **Supplementary Notes**, available upon publication.

## Code availability

The Python code for PepChem models, executable scripts, including training and benchmarks, and their usage instructions will be made available in an open source GitHub repository upon publication of the manuscript.

## Data availability

The data will be made available upon publication.

## Author contributions

M.S., D.B.G., and R.J.X. conceived the study. A.D., D.K., and M.S. designed the machine learning models, with input from H.L. R.J.X. supervised all aspects of the study. A.D. implemented the machine learning models and generated predictions for validation experiments. A.D. and D.K. performed statistical data analyses. A.M.B., A.J., and D.B.G. designed and performed *in vitro* peptide binding assays. J.G.A., K.R.C., and G.M.L. measured, analyzed, and curated the mass spectrometry peptidomics datasets. A.D., M.S., D.K., H.L., D.B.G., and R.J.X. wrote the manuscript. All authors provided feedback on the manuscript.

## Acknowledgements

The authors thank Theresa Reimels for editorial and graphic design support. We thank Steven A. Carr (Broad Institute) for facilitating analysis of proteomics datasets.

## Extended data figures

**Extended Data Figure 1:**
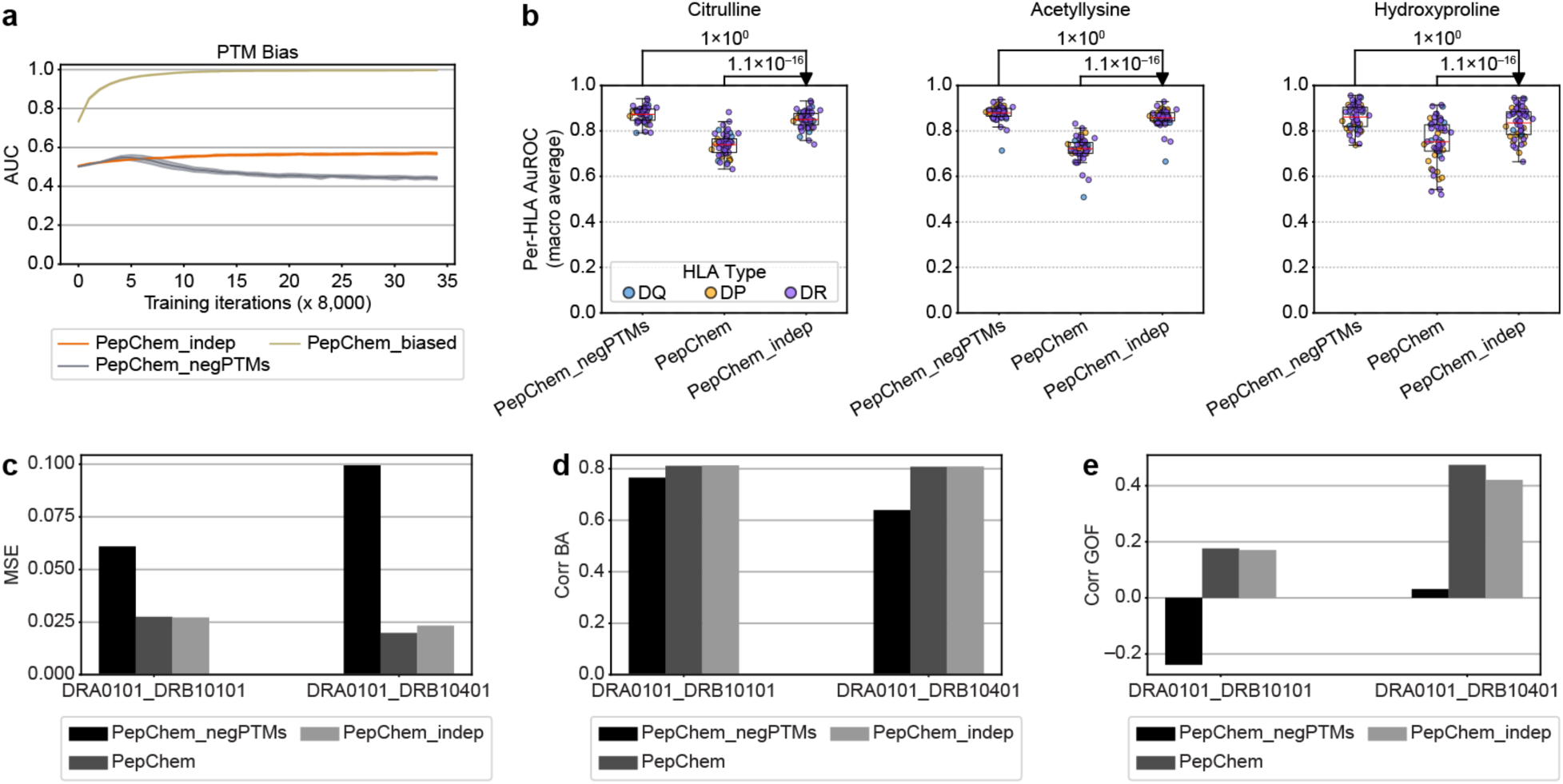
The artificial negative PTM set results in sub-optimal PTM representations and worsens citrulline BA predictions. **a** Bias test, measuring how likely a random PTM mutation is to increase the predicted EL presentation probability. **b** Performance on separation between positive PTM-peptides and negative standard amino acid peptides. **c** Mean squared error between predicted and true BA on citrullinated peptides (on D*_Sydney_* data). **d** Linear correlation coefficients between predicted and true BA values for citrullinated peptides (on D*_Sydney_* data). **e** Linear correlation coefficients for the predicted BA difference between arginine and citrulline form peptides (on D*_Sydney_* data).

**Extended Data Figure 2:**
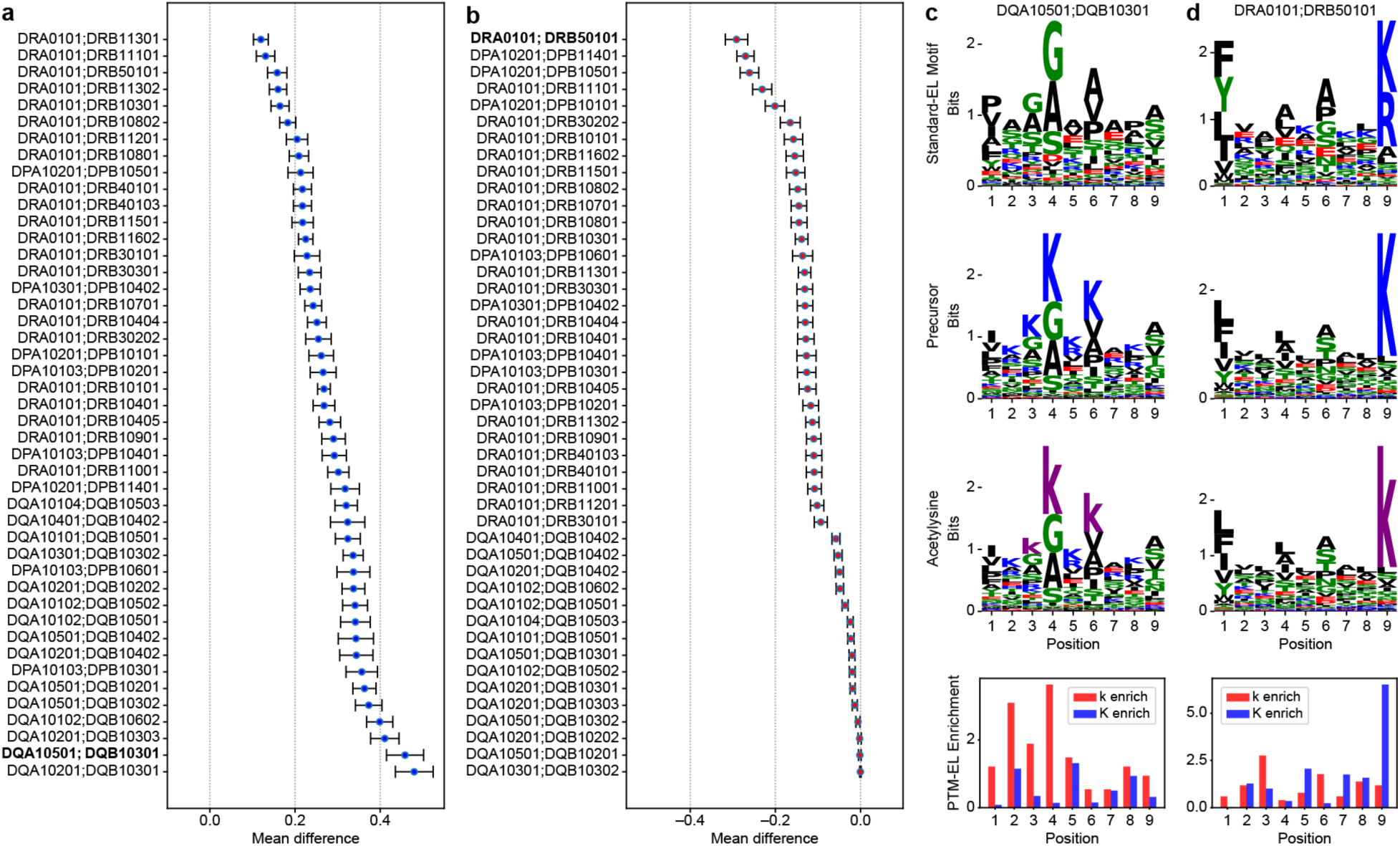
PepChem showcases preferential acetyllysine-induced gain and loss of function in peptide-HLA II interactions. **a** BA change distribution of 1,000 peptide pairs resulting in the largest gain of function from acetylation of lysine. **b** BA change distribution of 1,000 peptide pairs resulting in the largest loss of function from acetylation of lysine. **c** Gain-of-function analysis for DQA1*05:01;DQB1*03:01. From top to bottom: *i*) DQA1*05:01;DQB1*03:01 core extracted from the EL validation data of D*_net_*; *ii*) precursor motif that, upon acetylation of lysine, incurs the largest gain of function; *iii*) resulting modified motif that incurs a large gain of function upon lysine acetylation; and *iv*) increased frequencies of the PTM and its precursor residue at various positions based on the D*_PTM_*_–*EL*_ data, for the high gain- and loss-of-function alleles. **d** Loss of function analysis for DRB5*01:01.

**Extended Data Figure 3:**
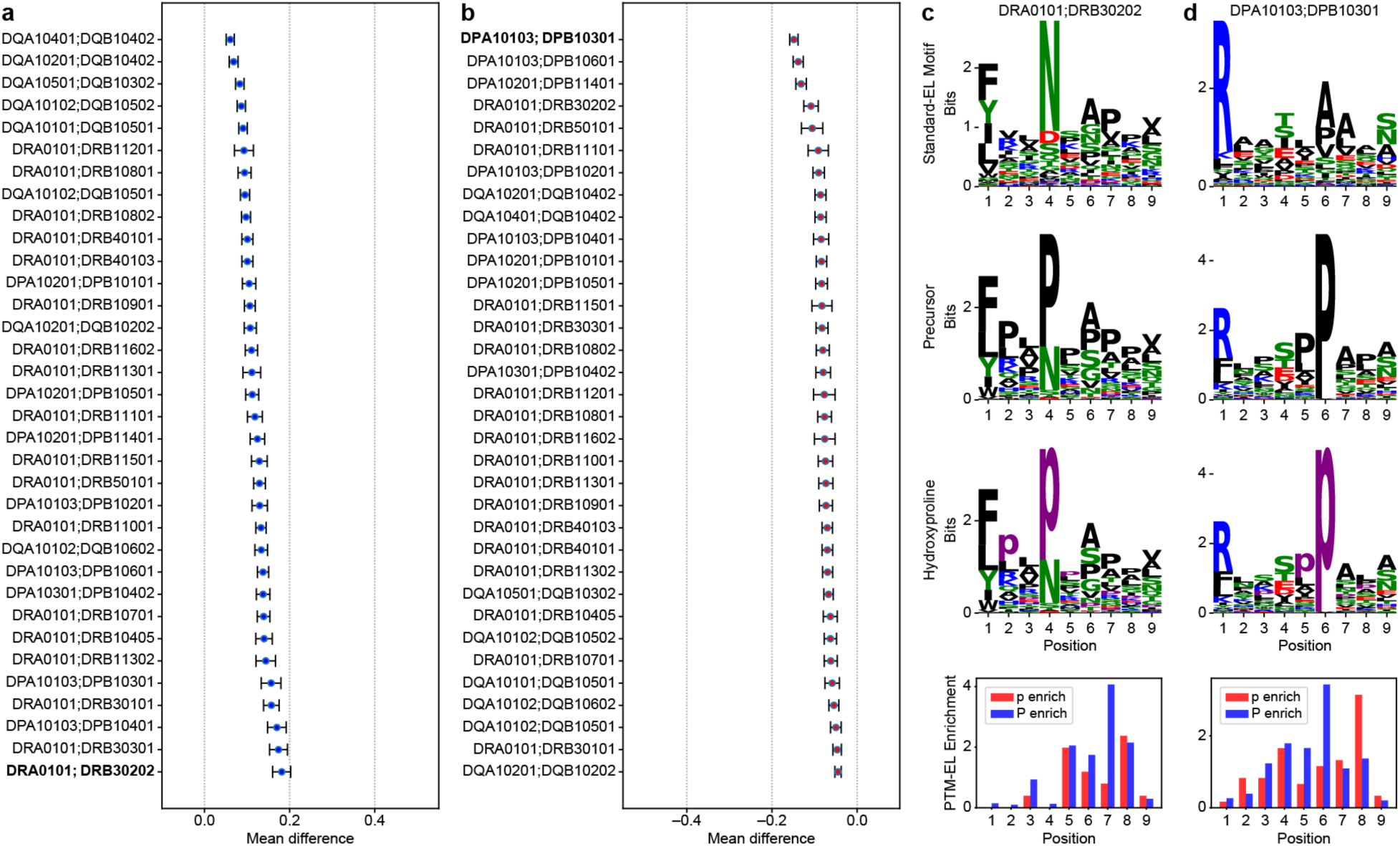
PepChem showcases preferential hydroxyproline-induced gain and loss of function in peptide-HLA II interactions. **a** BA change distribution of 1,000 peptide pairs resulting in the largest gain of function from hydroxylation of proline. **b** BA change distribution of 1,000 peptide pairs resulting in the largest loss of function from hydroxylation of proline. **c** Gain-of-function analysis for DRB3*02:02. From top to bottom: *i*) DRB3*02:02 core extracted from the EL validation data of D*_net_*; *ii*) precursor motif that, upon proline hydroxylation, incurs the largest gain of function; *iii*) resulting modified motif that incurs a large gain of function upon proline hydroxylation; and *iv*) increased frequencies of the PTM and its precursor residue at various positions based on the D*_PTM_*_–*EL*_ data, for the high gain- and loss-of-function alleles. **d** Loss of function analysis for DPA01*01:03;DPB1*03:01.

